# Crude *Necator americanus* worm extract diminishes pancreatic islets destruction in diabetic non-obese mice (NOD)

**DOI:** 10.1101/2020.03.03.975953

**Authors:** Teresiama Velikkakam, Bruna Garzedim de Araujo, Érica Alessandra Rocha Alves, Ricardo Toshio Fujiwara, Lilian Lacerda Bueno, Jacqueline Araújo Fiuza, Soraya Gaze

## Abstract

Hygiene hypothesis dictates that the lack of microbial interaction during the first childhood increases the chance of developing autoimmune diseases due to not proper immune system maturation. Helminthes are known by their Th2 and modulatory immune response induction. Here, it was evaluated the influence of *Necator americanus* antigens during type 1 experimental mouse model (non-obese diabetic – NOD). Intraperitoneal injections for 18 weeks did not impair an inflammatory response, but induced a mixed Th1/Th2 response with presence of IL4 and IL10 from different sources. However, the induced immune response was not sufficient to decrease glucose levels but showed a change in the inflammatory infiltrate in the pancreas. It is necessary more refined studies to clarify the mechanisms of how *Necator americanus* could impair diabetes progression in mice.

## Introduction

Diabetes melittus type 1 (DM1) is a disease characterized by pancreatic β cell selective destruction due to an autoimmune response (KAHALY; HANSEN, 2016). Its pathogenesis has many components, and in humans involves dysfunction in 40 major histocompatibity complex loci (HLA) (ATKINSON, 2014). Environment elements, like diet and infections, have an important role in this disease progress since they affect the immune system, especially in genetic susceptible people (CLARK; KROGER; TISCH, 2017). In the majority of the cases, DM1 is latent and asymptomatic for several years. Clinical manifestations only appear if 60-90% of pancreatic β cells are destroyed by the immune system and insulin production is not enough to control glucose levels in the blood (COPPIETERS; VON HERRATH, 2009).

It is known that the destruction of β cells is mediated by T and B cells, NK cells and APCs infiltrated in the islets of Langerhans (RICHARDSON; MORGAN; FOULIS, 2014; ZÓKA; SOMOGYI; FIRNEISZ, 2012). Several studies have demonstrated that, during the progression of the disease, the activation of T regulatory (Treg) cells culminates with a modulatory response capable of stopping the β cell autoantibodies production and autorreactive T cells effectiveness, evolving in a downmodulation in DM1 (ASKENASY, 2016; ATKINSON; EISENBARTH; MICHELS, 2014; CLARK; KROGER; TISCH, 2017; FERRETTI; LA CAVA, 2016).

Helminth infections are described as modulators of inflammatory responses due to their capacity of strong induction of Th2 and Treg immune responses (CANÇADO; FIUZA; DE PAIVA; LEMOS *et al.*, 2011; MOTRAN; SILVANE; CHIAPELLO; THEUMER *et al.*, 2018). Some studies have shown that DM1 in NOD mice may be ameliorated by helminthes. This property have been attributed to the fact that epitopes from helminthes were able to change the microenviroment of DM1 inflammation inducing Treg cells to produce IL-4, IL-10 and TGF-β (BERBUDI; AJENDRA; WARDANI; HOERAUF *et al.*, 2016; EGRO, 2013). *Schistosoma mansoni* antigens secreted by eggs were also able to reduce Treg cell proliferation, preventing DM1 in 4 weeks old NOD mice (ZACCONE; BURTON; MILLER; JONES *et al.*, 2009). Other autoimmune diseases can also be moderated by helminthes infections in experimental models. Studies with autoimmune encephalomielitys (GRUDEN-MOVSESIJAN; ILIC; MOSTARICA-STOJKOVIC; STOSIC-GRUJICIC *et al.*, 2010), colitis (SMITH; MANGAN; WALSH; FALLON *et al.*, 2007), sclerosis (FLEMING, 2013) and rheumatoid arthritis (OSADA; SHIMIZU; KUMAGAI; YAMADA *et al.*, 2009) also showed that helminthes and/or their products were able to prevent or suppress autoimmune diseases. However, it is not clear which helminthes can ameliorate and how they act in experimental DM1. Thus, this study evaluated the effect of crude antigens from adult *Necator americanus* in NOD mice model of DM1. The therapeutic properties of this parasite have been widely investigated in other autoimmune diseases, as well as celiac disease in humans.

## Methods

### Mice

Non-obese diabetic (NOD) female mice with 4 weeks obtained from Science and Technology in Biomodels Institute (Fiocruz) were tested negative for helminthes and protozoa. They were kept for two weeks in the same environmental conditions used in the experimental protocol for adaptation. Animals were kept in micro-isolater racks during all experimental period All the manipulation was under a sterile environment. Mice were divided in control (PBS) and AdEx (treatment) with 15 mice in each study group . All experimental protocols were reviewed and approved by Federal University of Minas Gerais Ethical Committee (license number 389/2012).

### *Necator americanus* crude extract (AdEx)

Hamsters were infected, via gavage, with 100 *Necator americanus* infective larvae stage. At day 42 post infection, animals were euthanized and adult worms were removed from intestine, washed with PBS plus antibiotics and sonicated in PBS. The solution was centrifuged at 500 xg and supernatant was filtered at 0.22 μM. Protein concentration was measured using microBCA kit accoding to manufature’s protocol (Pierce). The antigen was kept at −80°C until used. Antigens from *N. americanus* were kindly donated by Dr Ricardo Fujiwara (UFMG).

### Experimental design

NOD female mice (6 weeks old) were individually identified and monitored for 18 weeks. During this period , groups received, 3 times a week, 100 μL of PBS (control) or 15 μg AdEx per dose by intraperitoneal injection. Once a week, tail blood samples were collected to manufacture blood smear slides and to measure glicemia. At the end of 18 weeks treatment, mice were euthanized with overdose of xylazine and ketamine injected in the peritoneal cavity. Blood, pancreas and spleen were collected to measure immunological parameters.

### Blood glicemia measurement

Once a week, tail blood was placed in a tape to measure glicemia (ACCU-Check Active, Roche). All collections were done around the same hour in the day to decrease variation. Blood glicemia was considered non-fasten since the food was available all the time. Values were expressed in mg/dL and levels higher than 200mg/dL were considered abnormal.

### Blood smear slides

Once a week, a drop was collected from the tail to make blood smear slides. Then, blood cells were stained with hematology staining kit (Laborclin), in accordance with manufacturer’s instructions. Cells were analyzed under optical microscopy and the relative number (%) of circulating eosinophils, lymphocytes, neutrophils, basophils and monocytes was assessed by counting one hundred leukocytes, following Vanilda and Nascimento protocol (Vanilda and Nascimento, 2014).

### Histological analysis

Pancreas was carefully removed, identified and placed in 4% formalin buffer for fixation. Briefly, H&E staining was performed on 4-μm sequential sections of formalin-fixed, paraffin-embedded after placed in sylanized glass slides using hematoxylin and eosin (ALVARENGA; SILVA; FIUZA; GAZE *et al.*, 2018). Slides were analysed using optical microscope and the most preserved cut was selected to count and analyze all the pancreatic islets per animal. Islets were classified according to morphology and inflammatory infiltrate: without insulitis; moderate insulitis, peri-insulitis, severe insulitis, according described before (UKAH; CATTIN-ROY; CHEN; MILLER *et al.*, 2017)

### Flow cytometry

After euthanasia, spleens were collected and macerated in PBS through a 70 μM cell strainer (BD Falcon). Cell suspension was centrifuged at 300 xg and supernatant was discarded. Red blood cells were lysed with sterile distilled water, and leucocytes were centrifuged at 300 xg, ressupended and counted in RPMI1640 (Gibco) suplemented with 10% heat inactivated fetal bovine serum (Cultilab) with gentamicina 40 μg/mL (Sigma-Aldrich). Leucocytes at 1×106 were then kept in short period culture (4 h) with Brefeldin A (10 μg) (Sigma-Aldrich), 5 μg PMA (Sigma-Aldrich) and 50 μg Inomycin (Sigma-Aldrich). Later, cells were centrifuged, Fc receptor was blocked according to manufacture’s protocol (BD Biosciences) and stained with surface and cytokines antibodies according Fiuza et al., 2015 (Vaccine 2015 (33) 208-288). Antibodies (eBiosciences) used were: CD3 FITC(clone 145–2C11), CD4 PerCP-Cy5.5(clone RM4-4), CD8 BV421(clone 53-6.7), CD62L BV605(clone MEL-14), CD25 BV510(clone PC61), Cells were then fixed, permeabilized and stained for the cytokines IL-4(clone 11B11), IL-12p40(clone C15.6), IFN-γ (clone XMG1.2), TNF-α (clone MP6-XT22), IL-5 (clone TRFK5), IL-17A (clone TC11-18H10.1) and IL-10 (clone JES5–16E3), all PE. Data for 1×105 lymphocytes (gated by forward and side scatter) were acquired with a FACSFortessa flow cytometer and analyzed using FlowJo software (both from BD Biosciences). Isotype controls were used in all experiments.

### Nitric oxide measurement

Nitric oxide was measured in the splenocytes 4 h culture’s supernatant and in the serum collected after 18 weeks treatment. It was used Griess indirect reaction according to (GRANGER; TAINTOR; BOOCKVAR; HIBBS, 1996). Plates were read in Spectra-Max® Plus384 microplate reader (Molecular Devices) at 540nm. Sample concentration was interpolated in a standard curve, using the sample diluents, and showed as μg/mL.

### Serum cytokine concentration

Blood was centrifuged at 10,000 xg and sera were obtained and stored at −80°C until individual testing for cytokine levels. Sera were thawed only once and serum levels of IL-17A, TNF-α, IL-6, IFN-α, IL-2, IL-4 and IL-10 were quantified using the BD Cytometric Bead Array (CBA) Mouse Th1/Th2/Th17 Cytokine kit (BD Biosciences) following the manufacturer’s instructions. In addition, serum levels of IL-6, IL-10, MCP-1, IFN-γ, TNF-α e IL-12p70 were measured using BD Cytometric Bead Array (CBA) Mouse Inflammation kit (BD Bioscienses) according to manufecturer’s protocol. Data were acquired with FACSVerse (BD Biosciences) and analyzed using Flowjo 6.0 (BD Biosciences). Liophilized cytokines were used as standard and values were expressed/showed as as pg/mL. All values below detection limit were considered zero.

### Real-time quantitative PCR

Fresh prepared splenocytes were centrifuged and supernatant was discarded. Trizol (Invitrogen) solution was added and RNA extraction was prepared according to manufacturer’s protocol and concentration measured by Nanodrop (Thermo Fisher). After treatment of the RNA with DNAse (RQI Promega,),cDNA was synthesizedusing SuperScript II (Invitrogen). For real-time PCR, 10ng of cDNA was used per reaction with Syber Green (Promega). All the primers (described in Table 1) were designed and optimized. The experiments were perfomed using Step One Plus equipment and results analysed by Step One Software (Applied Biosystems). Results are expressed by 2^^^(CT test- CT control), and beta-actin was chosen as housekeeping gene.

**Table 1:**
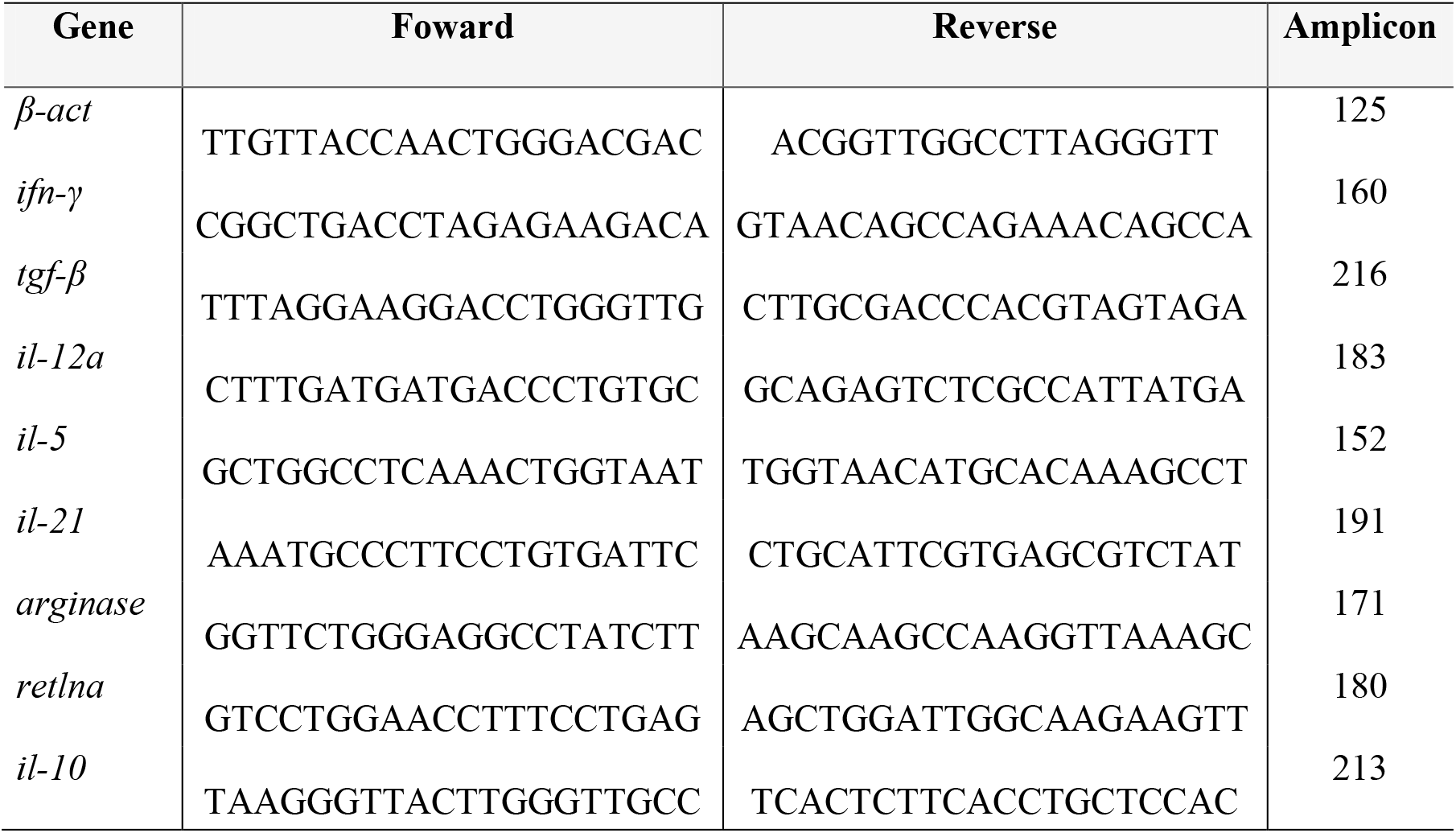
Primers designed for each tested gene used in real-time PCR.

### Statistical analysis

T test with Holm-Sidak post-test was used for multicomparisons. Also, two non-parametric tests Mann-Whitney test and Spearman correlation, according to analyzed data. Statistical significance was considered if p value ≤ 0.05. All data was analyzed using GraphPad Prism 6.0 software (GraphPad).

## Results

### Blood glicemia was not affected by AdEx treatment

Blood glucose levels were measured weekly, in the morning, for 18 weeks. During all experimental period, there was no difference in the circulating levels of glucose between AdEx-treated and placebo groups show (Figure 1A). During the 18 weeks treatment, four animals per control group and three mice per AdEx group showed levels above 600mg/dL and all died before the next measurement. Since all NOD mice were individually identified, glucose levels were also analyzed considering the first measurement (week 1). Results were plotted in heatmap (Figure 1B), where gray indicates lower variation and red higher variation compared to the first week of treatment. As demonstrated in the figure 1B, AdEx-treated group showed a tendency to have lower variation in the blood glucose levels than PBS group (Figure 1B).

**Figure 1.**
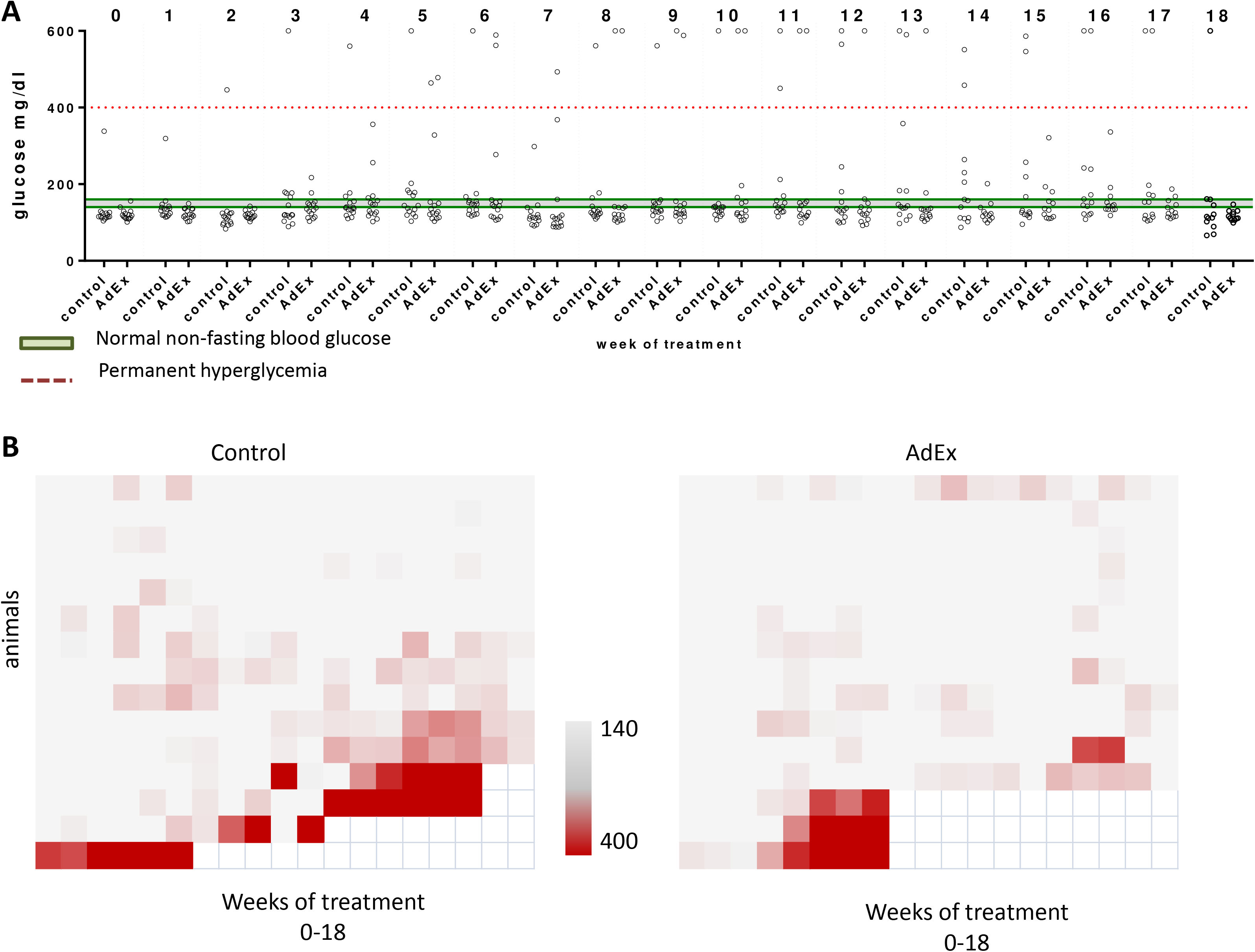
Blood glucose levels distribution after 18 weeks of AdEx treatment. **(A)** Blood glucose levels (mg/dL) in mice from treated (AdEx) and control (PBS) groups from week 0 to 18. Dashed line indicates level of permanent glycemia and gray area normal non-fasting glycemia levels according to (LEITER; PROCHAZKA; COLEMAN, 1987) **(B)** Heat map of glycemia levels per week per group. Lower level considered 140mg/dL (gray) and upper level 400 mg/dL (red). Empty spots indicated lost of measurement since mice did not survive.

### NOD mice presented decreased severity of insulitis after AdEx treatment

Pancreas from mice were evaluated by optical microscopy after H&E staining. The severity of insulitis was analyzed by the**_**total number of Langerhans islets and size of inflammation around them. When data was analyzed, AdEx group showed higher number of Langerhans islets than PBS group (Figure 2A). Moreover, Langerhans islets were larger than those found in the control group. The results also demonstrated that mice treated with AdEx showed 42.3% of islets without insulitis 11.5% with periinsulitis, 15.4% with moderate insulitis and 30.8% com severe insulitis. On the other hand, while animals treated with PBS showed 29.7% of islets without insulitis, 18.5% with periinsulitis, 11.1% with moderate insulitis, and 40.7% with severe insulitis (Figure 2B), indicating a decrease of the inflammatory response in the AdEX-treated group.

**Figure 2.**
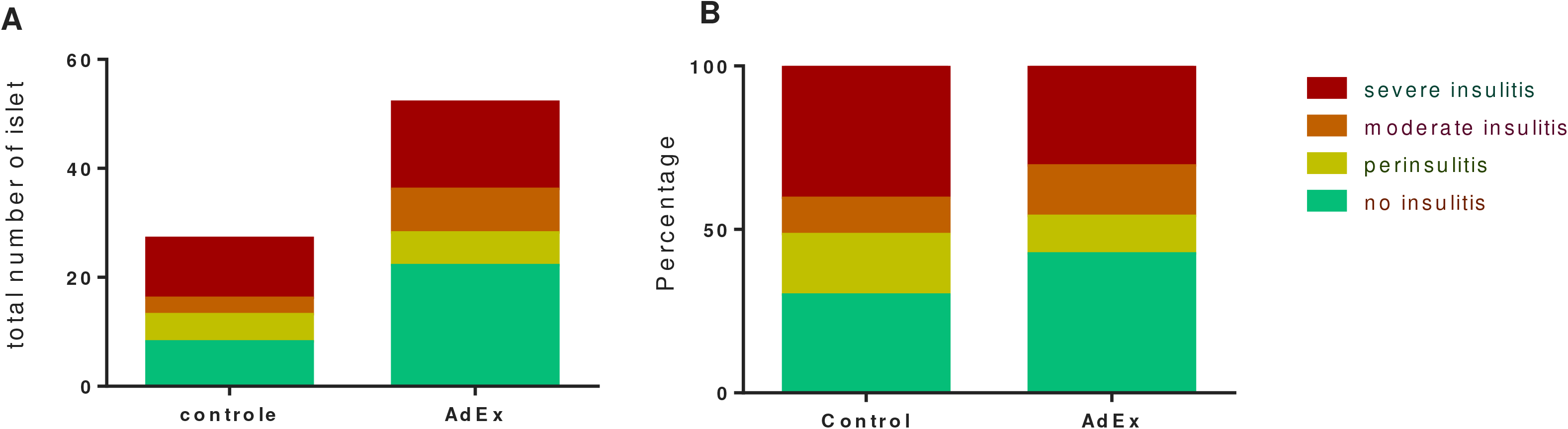
Evaluation of inflammatory infiltrate in Langerhans islets in NOD mice treated with AdEx. Total number (A) and percentage (B) of Langerhans islets presenting or not inflammatory infiltrate.. From bottom to top: green – no insulitis, yellow – perinsulitis, orange – moderate insulitis, red – severe insulitis.

### AdEx treatment induces an increase of circulating eosinophils

AdEx injections increased the relative number of circulating eosinophils. This increase was significant as early as second week of treatment and was kept elevated through the whole treatment compared to the PBS control group (Figure 3). The relative number of lymphocytes was also increased in the AdEx group (data not shown). The other cell subtypes analyzed, including neutrophils, mastocytes, basophils and monocytes, showed no difference between both analyzed groups.

**Figure 3.**
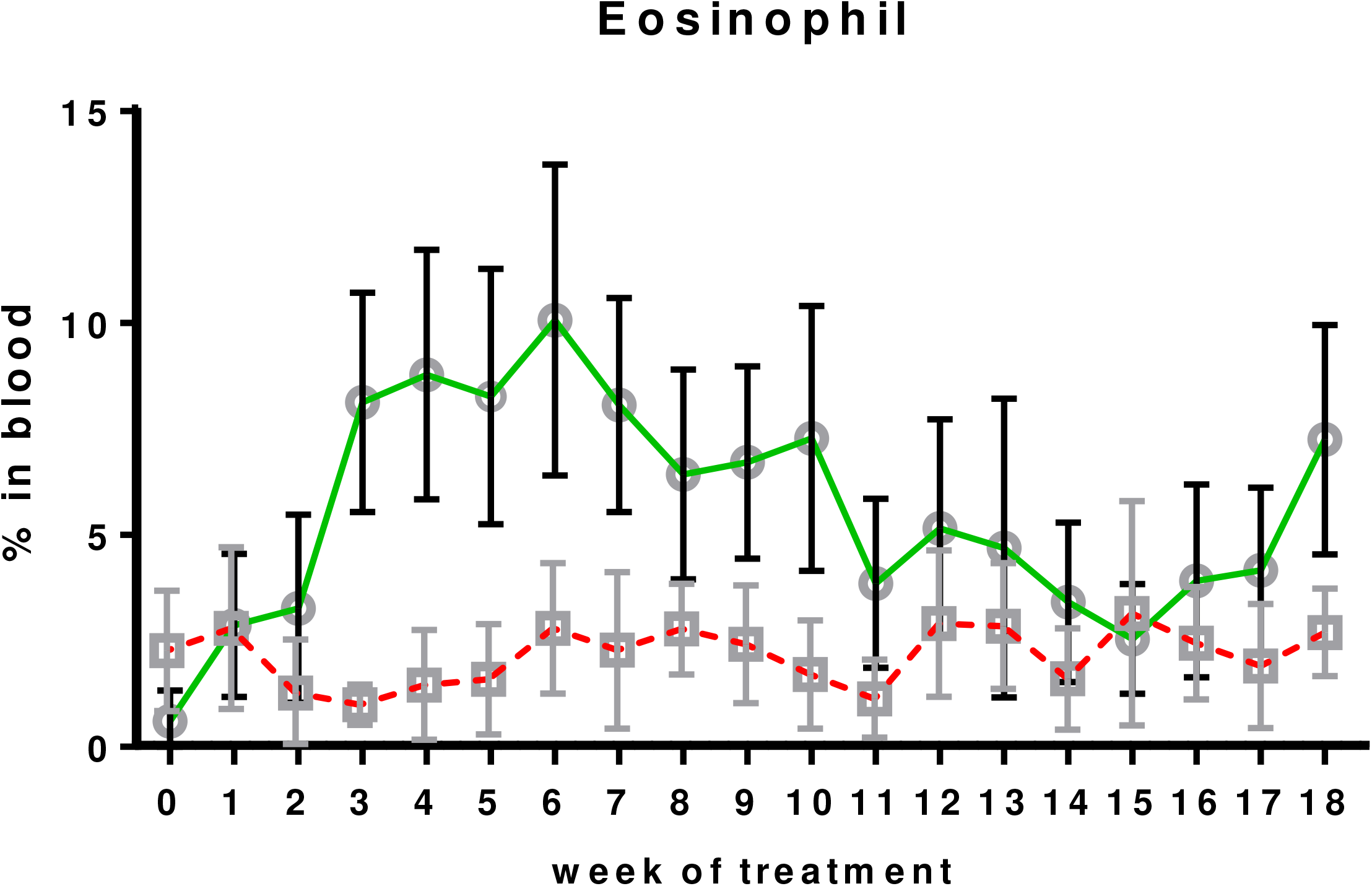
Percentage of eosinophils in the blood circulation. Square – control group, circle AdEx group. Data presented as mean ± SEM.

### A mixed immune response profile was observed after AdEx treatment

The profile of cytokines induced by AdEx treatment was evaluated in two compartments: Sera and splenocytes. In the sera from NOD mice, the AdEx treatment induced an increase of IL-2 (p=0.0138; Figure 4A), IFN-γ (p=0.0072; Figure 4B), IL-12 (p=0.0138; Figure 4C) and IL-6 (p=0.0099; Figure 4D), all considered inflammatory cytokines, compared to the control group. In addition, AdEx-treated mice presented increased serum levels of IL-4 (p=0.0009; Figure 4E) and IL-10 (p=0.0138; Figure 4F) compared to the PBS-treated animals, showing that the AdEx treatment induced a mixed profile of circulating cytokines. Since we have observed increased production of modulatory cytokines in sera from AdEx-treated mice, we also analyzed short cultured splenocytes, in order to know which cells were producing those cytokines. Interesting, our results demonstrated that, after 18 weeks treatment, the AdEx group have increased IL-4 production by CD4^+^ (p=0.0086; figure 5A) and CD8^+^ (p=0.0129; figure 5B) T cells. . In addition, AdEx mice presented higher IL-10 production, not only by CD4^+^ (p=0.0086; figure 5C) and CD8^+^ (p=0.0086; figure 5D) T cells, as well as by macrophages (p=0.0086; figure 5F), compared to the PBS group. Furthermore, in the splenocytes culture from AdEx group, the production of IL-10 was also higher by the subpopulation of CD4^+^ T cells expressing CD25 (CD4^+^CD25^+^) than in the culture from PBS control group (p=0.031, figure 5E). Taken together, results from splenocytes culture showed that treatment with ADEX increased the production of modulatory cytokines by splenic leukocytes. Other cytokines measured according to described before did not show statistical differences between analyzed groups.

**Figure 4.**
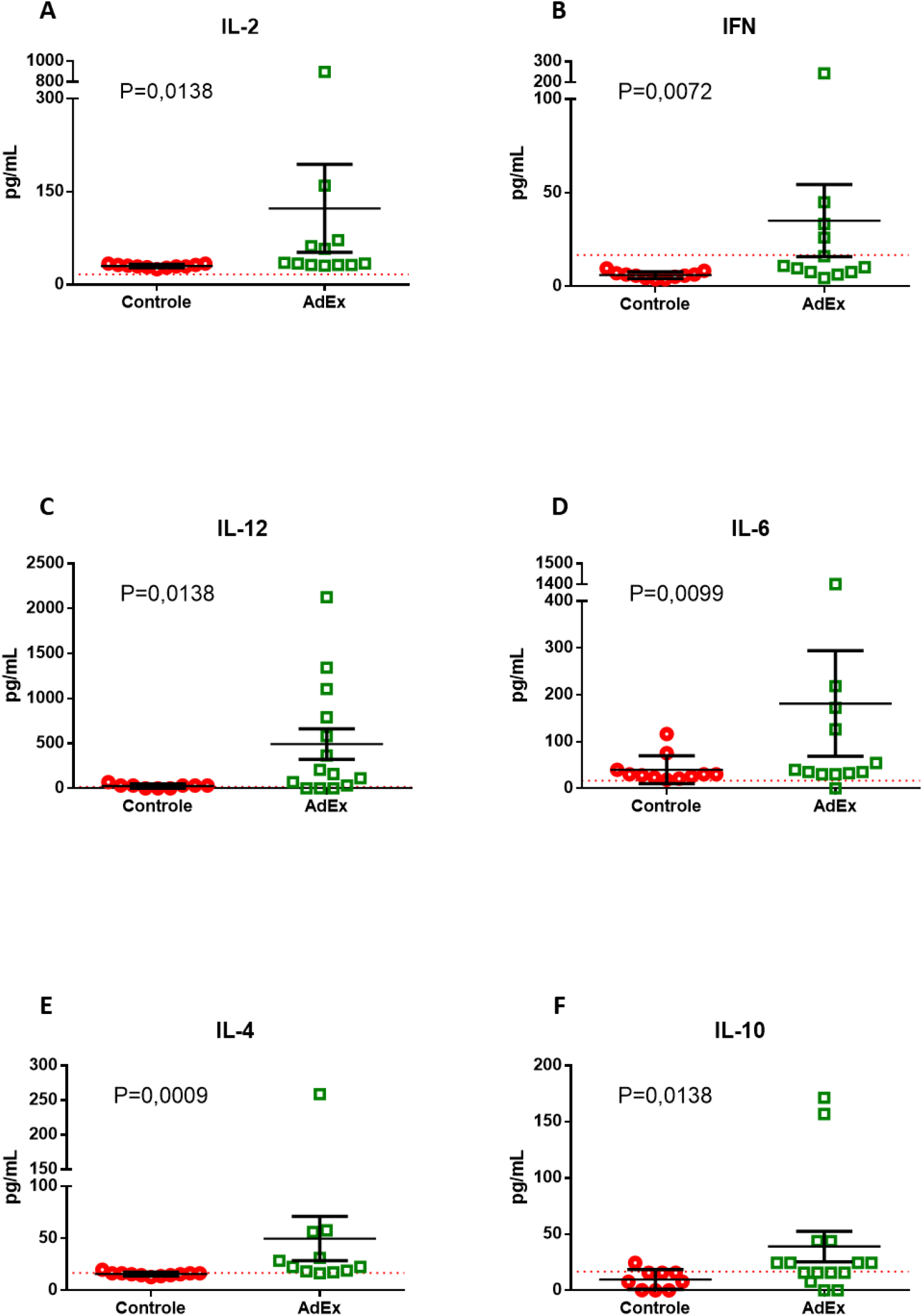
Serum levels of cytokines in NOD mice treated or not with AdEx. . (A) IL-2; (B) IFN-γ; (C) IL-12; (D) IL-6; (E) IL-4; (F) IL-10. Red circles – control (PBS) group. Green square – treated (AdEx) group. Statistical analysis was performed using unpaired T test. Data presented as mean ± SEM. Dashed line indicates detectable levels according to manufacture’s protocol.

**Figure 5.**
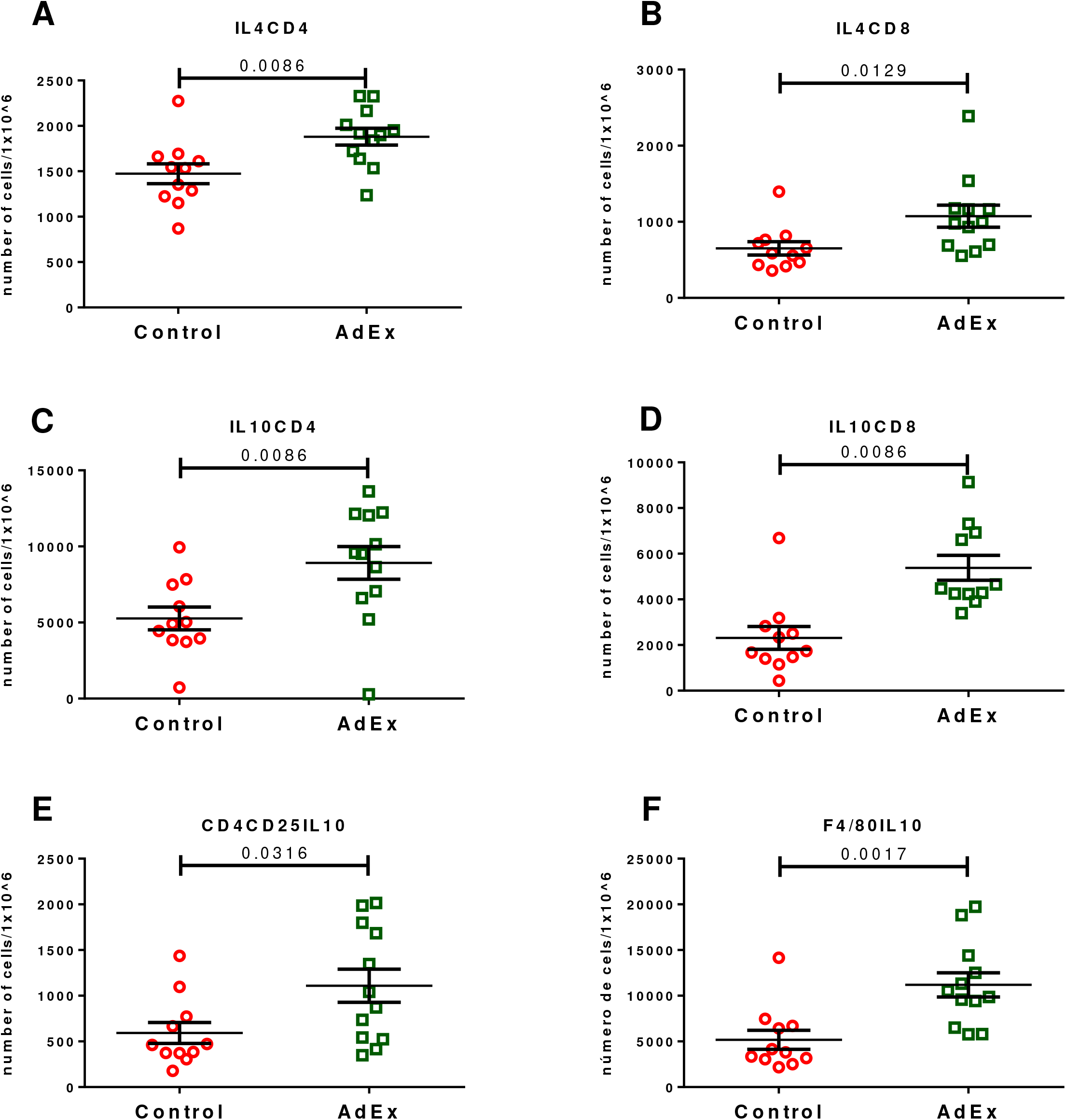
Evaluation of the cytokine levels in the culture supernatant of splenocytes from NOD mice treated or not with AdEx. (A) IL4CD4; (B) IL-4CD8; (C) IL10CD4; (D) IL10CD8; (E) CD4CD25IL-10; (F) F4/80IL-10. Red circles – control (PBS) group. Green square – treated (AdEx) group. Statistical analysis was performed using unpaired T test. Data presented as mean ± SEM.

Levels of mRNA related with inflammatory and modulatory immunological profiles were evaluated (IFN-γ, IL-12a, IL-5, IL-21, Retnla, Arginase, TGF-β and IL-10) were analyzed in splenocytes by qPCR. RETNLA (Fizz1) showed higher expression in AdEx group (p=0.0116) compared to control (Figure 6A). However, the expression of other regulatory genes, as Arginase, TGF-β and IL-10, was lower in AdEx mice compared to control group (p=0.0117, p= 0.0369 and p=0.0113, respectively) (Figure 6B, C and D, respectively). No statiscal differences were observed when analyzing the expression of IFN-γ, IL-12a, IL-5, IL-21 (data not shown).

**Figure 6.**
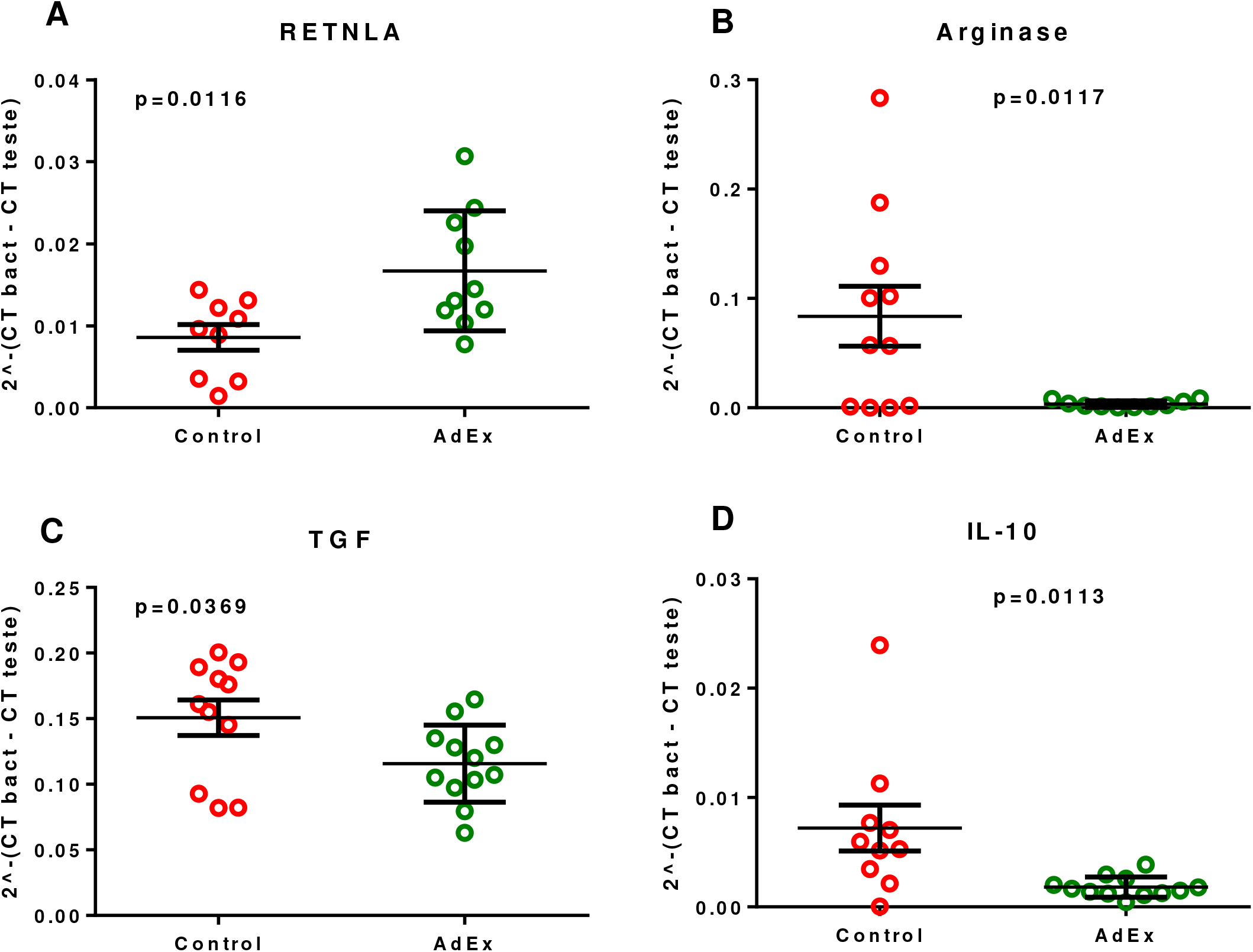
Fresh spleen cell mRNA expression evaluation. (A) RETNLA (FIZZ-1); (B) Arginase; (C) TGF; (D) IL10. Red circles – control (PBS) group. Green square – treated (AdEx) group. Statistical analysis was performed using unpaired T test. Data presented as mean ± SEM.

### AdEx treatment induces decrease of Nitric Oxide

The levels of NO were analyzed in sera and culture supernatant from splenocytes by Griess reaction. In both, the treatment with AdEx for 18 weeks induced lower levels of nitric oxide when compared to control PBS group (sera, p=0.002 and supernatant, p=0.003) (Figures 7A and B), showing that AdEx injection is influencing free radicals production in NOD mice..

**Figure 7.**
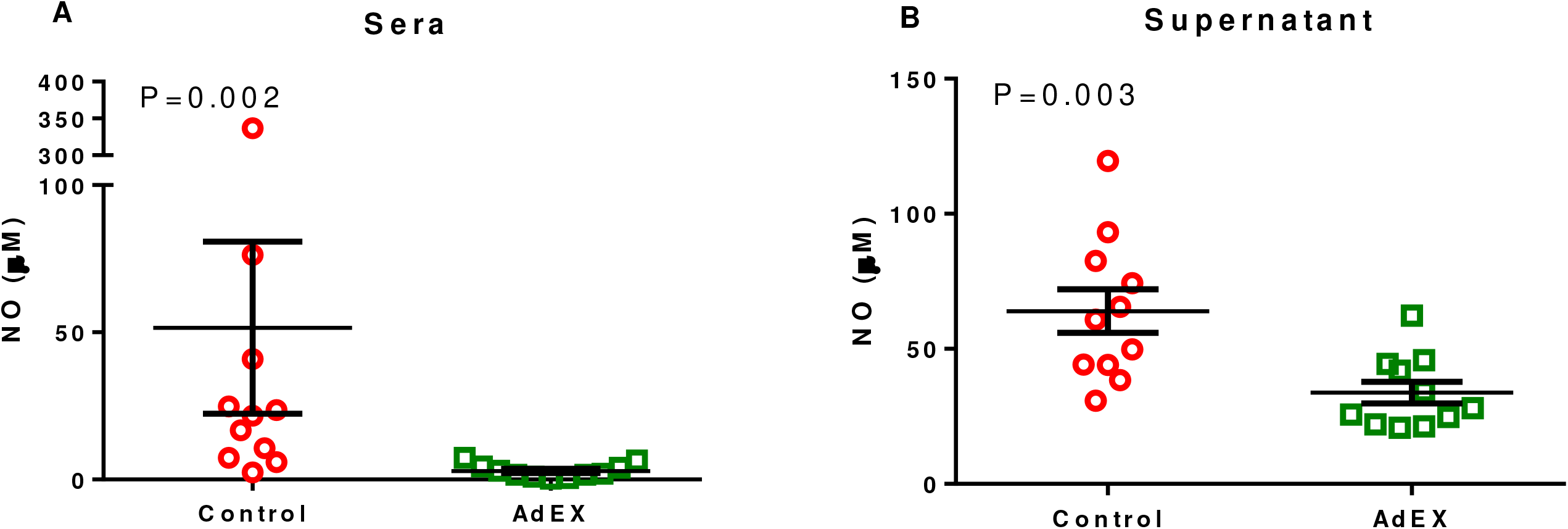
Nitric oxide production evaluation. (A) Sera; (B) Supernatant of 4h spleen cell culture. Red circles – control (PBS) group. Green square – treated (AdEx) group. Statistical analysis was performed using unpaired T test. Data presented as mean ± SEM.

### Higher levels of glucose is negative correlated to levels of modulatory cytokines

Correlations were made between levels of glucose and production of IL-4 and IL-10 by CD4^+^ and CD8^+^ T cells. After 18 weeks of treatment with AdEx, mice that presented higher levels of glucose showed a decrease of IL4CD4 (p=0.0183, r=0.6714; figure 8A), IL4CD8 (p=0.0174, r=0.6749; figure 8B) and IL10CD4 (p=0.00264, r=0.6797; figure 8C), being correlated negatively.

**Figure 8.**
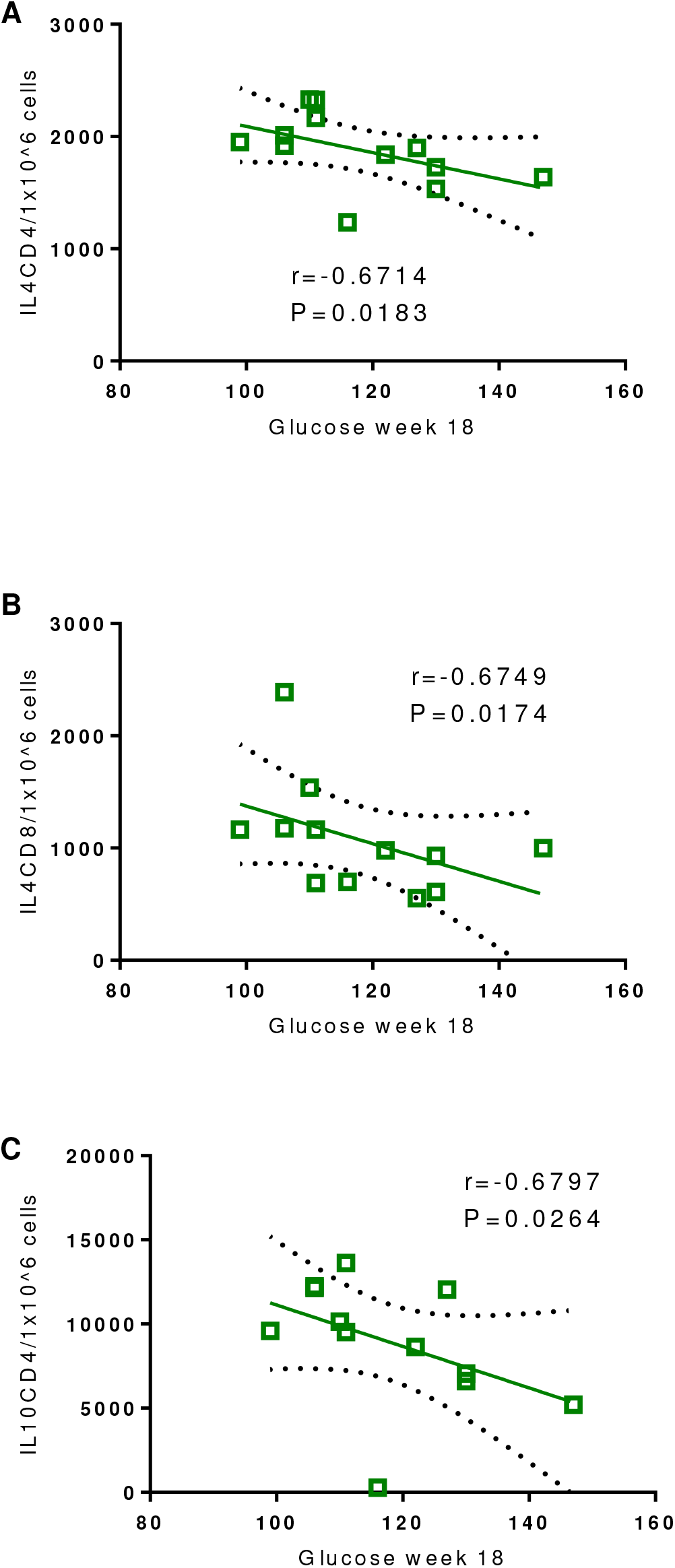
Correlation between 4h culture of spleen cell population and glucose level at week 18. (A) IL4CD4; (B) IL-4CD8; (C) IL10CD4 versus glucose level at week 18 in treated (AdEx) group (green square). Green line correlation. Dashed line: 95% of distribution.

## Discussion

The use of helminthic parasites as therapeutic tools for chronic diseases has been studied for many groups (CANÇADO; FIUZA; DE PAIVA; LEMOS *et al.*, 2011; CROESE; O’NEIL; MASSON; COOKE *et al.*, 2006; ELLIOTT; LI; BLUM; METWALI *et al.*, 2003; ELLIOTT; SETIAWAN; METWALI; BLUM *et al.*, 2004; ELLIOTT; URBAN JF; ARGO; WEINSTOCK, 2000; HÜBNER; SHI; TORRERO; MUELLER *et al.*, 2012; IMAI; TEZUKA; FUJITA, 2001; KHAN; BLENNERHASSET; VARGHESE; CHOWDHURY *et al.*, 2002; LIU; SUNDAR; MISHRA; MOUSAVI *et al.*, 2009; REARDON; SANCHEZ; HOGABOAM; MCKAY, 2001; SAUNDERS; RAINE; COOKE; LAWRENCE, 2007; SUMMERS, R. W.; ELLIOTT, D. E.; URBAN, J. F.; THOMPSON, R. *et al.*, 2005; SUMMERS, R. W.; ELLIOTT, D. E.; URBAN, J. F.; THOMPSON, R. A. *et al.*, 2005; SUTTON; ZHAO; MADDEN; ELFREY *et al.*, 2008). Particularly for DM1 , the use of pathogens or pathogen-derived products have shown to modulate autoimmune diabetes in the NOD mice (HARN; MCDONALD; ATOCHINA; DA’DARA, 2009; HÜBNER; SHI; TORRERO; MUELLER *et al.*, 2012; HÜBNER; STOCKER; MITRE, 2009; IMAI; TEZUKA; FUJITA, 2001; LIU; SUNDAR; MISHRA; MOUSAVI *et al.*, 2009; SAUNDERS; RAINE; COOKE; LAWRENCE, 2007; SUTTON; ZHAO; MADDEN; ELFREY *et al.*, 2008; ZACCONE; FEHÉRVÁRI; JONES; SIDOBRE *et al.*, 2003). The present work also demonstrated important implications in NOD mice, as a mixed Th1/Th2 response and the decrease of damage in the pancreas. It is important to note that in our study, mice showed lower incidence of diabetes than previously reported (Liu et al., 2009). It is known that the age is one important factor to influence the normal glucose levels. In NOD mice, there are evidences that diabetes may develop as late as 30 weeks of life (KAMINITZ; MIZRAHI; ASKENASY, 2014). Unfortunately, at the end of the experimental period, NOD mice were 24 weeks old because AdEx was available to perform treatment just for 18 weeks.

When mice were individually compared, there is a tendency of AdEx treatment to keep glucose levels near the week 0 levels. Interestingly, the glucose blood levels corroborate the inflammation found in the pancreas. Histology analyses showed that the pancreatic islet destruction was not different in numbers from AdEx to control group. Thus, could be a reason that diabetes had lower rates in this study. However, the AdEx treatment seems to influence the inflammatory infiltrate in the pancreas since 42% of the islets had no insulitis in treat group compared to 29.7% in control group.

It has been shown that *N. americanus* infection can induce a strong Th2 response, but Th1 component is also present (GEIGER; CALDAS; MC GLONE; CAMPI-AZEVEDO *et al.*, 2007). AdEx used in this study is composed of proteins from whole adult worms, males and females, and could explain the mixed response observed. Eosinophils have been shown to play an important effector role in helminth infections. They could immunopolarize the response (WEBB; TAIT WOJNO, 2017), inducing preferably Th2 response through releasing mediators like Indoleamine 2,3-dioxygenase and Eosinophil-derived neurotoxin (SPENCER; WELLER, 2010). Since there was an increase of eosinophils in early weeks of injection, they could contribute to favor a strong Th2 immune response, as observed, in the presence of *N. americanus* antigens and could be one of the mechanisms to decrease damage in the pancreas.

Splenocytes were also affected by AdEx treatment demonstrating the capacity to influence several immune sites. After 18 weeks treatment, CD4^+^ and CD8^+^ T cells producing IL-4 and IL-10 showed an increased numbers in the spleen compared to control placebo group. Other studies had shown that helminthes can stimulate Treg cells and other types inducing IL-10 and IL-4 production, and this could ameliorate inflammation caused by Th1 and Th17 (GAZE; DRIGUEZ; PEARSON; MENDES *et al.*, 2014; MCSORLEY; HEWITSON; MAIZELS, 2013; WEBB; TAIT WOJNO, 2017). This is particularly important since NOD mice have shown lymphopenia, especially for CD4^+^ and Treg cells (DEJACO; DUFTNER; GRUBECK-LOEBENSTEIN; SCHIRMER, 2006; KING; ILIC; KOELSCH; SARVETNICK, 2004). It was also described a lower IL-4 intrinsic production by NOD mice (AOKI; BORCHERS; RIDGWAY; KEEN *et al.*, 2005; DELOVITCH; SINGH, 1997). Thus, AdEx treatment increased the IL-4 and IL-10 not only by splenocytes but also blood circulating cytokines, another way to help controlling the inflammation in the pancreas.

Moreover, IL-12, IL-2 and IL-6 levels were increased in treated mice sera. Nevertheless, how those cytokines balance in different compartments are interfering in the DM1 pathology still needs a deeper understanding. The corroboration that the steady state of AdEx treatment, even if it is only for 18 weeks, is an attempt of controlling inflammation, also demonstrated by lower levels of nitric oxide production in treated group compared to control group. This could be a result of a possible alternative macrophage induction by AdEx producing IL-10. Alternative activated macrophages express Arginase-1 inducing L-ornitin production and are involved in wound healing (REYES; TERRAZAS, 2007). On the other hand, classic macrophages convert arginin in nitric oxide causing lesion by respiratory burst and β cells killing (REYES; TERRAZAS, 2007). This could be one of the factors involved in the pancreatic islets preservation observed in AdEx group. The induced mixed profile was also shown by mRNA levels. Although Arginase mRNA was decreased in AdEx treated group, while FIZZ1 (Retnla) was increased compared to control group. FIZZ1 is a molecule expressed by alternative macrophages and is also involved in wound healing (ANTHONY; RUTITZKY; URBAN; STADECKER *et al.*, 2007). Other studies also demonstrate that helminthes induce higher levels of FIZZ1 in mice (ANTHONY; URBAN; ALEM; HAMED *et al.*, 2006; NAIR; GALLAGHER; TAYLOR; LOKE *et al.*, 2005).

An association between AdEx treatment and diabetes amelioration were demonstrated by correlations between blood glucose levels and CD4^+^ and CD8^+^ T cells producing IL-4 and IL-10. Those negative correlations suggest a potential effect of AdEx to reduce the damage caused by diabetes type 1 in mice. Thus, this study corroborates the hypothesis of helminthes antigens can potentially modulate and modify the immune response to benefit mice affected by DM1. Other studies with infections or antigen administration using *Schistosoma mansoni, Taenia crassiceps , Heligmosomoides polygyrus, Trichinella spiralis ou Strongyloides venezuelensis* showed that helminthes parasites prevented or inhibited DM1 in NOD mice (AJENDRA; BERBUDI; HOERAUF; HÜBNER, 2016). However, this positive effect only happened if antigen administration or infection started before insulitis establishment, usually around 4 weeks old in NOD mice (BERBUDI; AJENDRA; WARDANI; HOERAUF *et al.*, 2016).

In conclusion, this study showed the potential therapeutic use of *Necator americanus* antigens to prevent DM1 progression by decreasing β cell destruction.This study supports the need for further analysis aiming at elucidating the helminth-induced mechanisms in DM1.

## Acknowledges

This project had financial support from Conselho Nacional de Pesquisas (CNPq) Universal grant number 444060/2014-6 and Rene Rachou Institute-Fiocruz MG. TV, BGA had scholarship, and JAF had fellowship, from CNPq. RTF and LLB have career fellowship from CNPq. Data and results provided by this manuscript are not plagiarism and were not published anywhere before here. The authors also thank the flow cytometry platform of the Fiocruz Program for the Technological Development Tools for Health (PDTIS). We are very thankful for the help of Dr Marcelo Antônio Pascoal Xavier.

## Author’s contribution

TV, BGA, EARA, JAF and SG participated in all experiments and analysis. TV, EARA, RTJ, LLB, JAF and SG wrote the manuscript. JAF, EARA and SG designed experiments. RTJ and LLB provided crude *N. americanus* extract.

